# Hypertonic saline and aprotinin based blockage of SARS-CoV-2 specific furin site cleavage by inhibition of nasal protease activity

**DOI:** 10.1101/2021.11.19.469276

**Authors:** Clarissa Li, Adam W. Li

**Affiliations:** EpigenTek Group Inc., 110 Bi County Boulevard, Suite 122, Farmingdale, NY, 11735, USA

## Abstract

SARS-CoV-2 enters into the human body mainly through the nasal epithelial cells. Cell entry of SARS-CoV-2 needs to be pre-activated by S1/S2 boundary furin motif cleavage by furin and/or relevant proteases. It is important to locally block SARS-CoV-2 S1/S2 site cleavage caused by furin and other relevant protease activity in the nasal cavity. We tested hypertonic saline and aprotinin-based blockage of SARS-CoV-2 specific furin site cleavage by furin, trypsin and nasal swab samples containing nasal proteases. Our results show that saline and aprotinin block SARS-Cov-2 specific furin site cleavage and that a saline and aprotinin combination could significantly reduce SARS-Cov-2 wild-type and P681R mutant furin site cleavage by inhibition of nasal protease activity.

## Introduction

The recent COVID-19 pandemic was caused by SARS-CoV-2, a new member of the same coronavirus family that caused SARS and MERS. To date, more than 256 million of infections and 5.1 million of deaths have been reported worldwide [1], and the numbers continue to rise because of increased CoVID-19 variants that result from viral mutations. There are no effective drugs that can be used for the prevention and treatment of COVID-19. The vaccines against CoVID-19 achieved marvelous effects on preventing/reducing original SARS-CoV2 infections. However, the vaccine effects are significantly reduced against CoVID-19 variants such as Delta variant that is the dominant one in the USA and many other countries currently [2, 3].

It was found that the SARS-CoV-2 spike (S) glycoprotein harbors a furin cleavage site (FCS) at the boundary between the S_1_/S_2_ subunits, which could be cleaved by furin and/or relevant proteases secreted from host cells. Unlike SARS-CoV, cell entry of SARS-CoV-2 needs to be pre-activated by furin and/or relevant proteases, reducing its dependence on target cell proteases for entry. The cleavage activation of S-protein is well demonstrated to be essential for SARS-CoV-2 spike-mediated viral binding to the host ACE2 receptor, cell-cell fusion, and viral entry into human lung cells [4, 5]. It was also observed that other viruses containing a furin cleavage site, such as H5N1, displayed increased replicates and developed higher pathogenicity [6].

The complete SARS-CoV-2 furin cleavage site has been characterized as a 20 amino acid motif that is corresponding to the amino acid sequence ^A^672-^S^691 of SARS-CoV-2 spike protein (QOS45029.1), with one core region SPRRAR│SV (eight amino acids, ^S^680-^V^687) and two flanking solvent accessible regions (eight amino acids, ^A^672–^N^679, and four amino acids, ^A^688-^S^691). The core region is very unique as its ^R^683 and ^A^684 positions are positively charged (Arg) and hydrophobic (Ala) residues, respectively, which allow this site to not only be cleaved by serine protease furin or furin-like PCs, but also permit cleavage efficiency to be facilitated by other serine proteases targeting mono- and dibasic amino acid sites such as matriptase, kallikrein 1 (KLK1), human airway trypsin (HAT), and TMPRSS2. Furthermore, mutation of ^P^681 (non-polar proline) to positive charge ^H^681 (histidine) or even more positively charged ^R^681 to form multibasic amino acid sites could increase S1/S2 boundary cleavage, thereby increasing viral replicates in human airway and transmission [7,8]. In addition, a serine at ^S^680 of the core region could also highly increase the cleavage efficiency, causing increased viral replication, unrestricted organ tropism, and virulence and mortality rates as proven in H5N1 infection studies in mice [9].

It was demonstrated that SARS-CoV-2 enters into the human body mainly through the nasal epithelial cells [10]. There are high levels of furin and other facilitating serine proteases, which are secreted from the nasal epithelial cells and generated from bacteria in the nasal cavity. Thus, it would be important to locally block SARS-CoV-2 S1/S2 site cleavage caused by furin and other facilitating serine protease activity in the nasal cavity as the furin-based SARS-CoV-2 S1/S2 cleavage increases SARS-CoV-2 entry into cells and its replication, and eventually develops higher pathogenicity and transmission of COVID-19. To address this, we tested saline and aprotinin-based blockage of SARS-CoV-2-specific furin site cleavage by furin and other facilitating serine proteases. Our results show that saline and aprotinin block SARS-Cov-2 specific furin site cleavage and that a saline and aprotinin combination could significantly reduce SARS-Cov-2 wild-type and P681R mutant furin site cleavage by inhibition of nasal protease activity.

## Materials and Methods

### Synthesis of peptide containing SARS-CoV-2 specific furin cleavage site

The dual-tagged peptides containing the wild-type, mutant (P681R), and triple R-deleted SARS-CoV-2-specific furin motif were synthesized through Biomatik platform Peptide Synthesis Services. The peptide is labeled with polyhistidine at N terminal and biotin at C terminal.

### Sample collection

Collection of nasal swab samples from healthy uninfected volunteers was in accordance with the standard CDC nasal swab collection protocol. The collected samples were released into 300 μl of protease cleavage (PC) assay buffer (EpiGentek) or NaCl solution at different concentrations by rotating the swab in the buffer for 30 sec. the solution was centrifuged at 12,000 rpm at RT for 1 min. The supernatant was collected and freshly used for assay.

### SARS-CoV-2 specific furin site cleavage measurement

The synthesized peptides containing the wild-type, mutant (P681R), and triple R-deleted SARS-CoV-2-specific furin motif were added at a concentration of 5 ng/well (prepared with PBS) to the blocked (2% BSA) Ni-NTA-coated strips and incubated for 1 h at RT. After washing the strips for 3 times with PBS-T, purified proprotein convertase furin (New England Biolabs) or protease trypsin (Sigma) was added at different concentrations and incubated for 25 min at 37°C. Protease cleavage (PC) assay buffer (EpiGentek) was used as the assay solutions. After washing for 4 times, streptavidin-HRP (100 μl, 1:5000 dilution in PBS-T) was added and incubated for 30 min at RT. After washing for 4 times, 100 μl of TMB solution were added per well and blue color development was monitored for 2-10 min. The reaction was stopped with an equal volume of 1 M HCl and the optical density was measured with a microplate reader at a wavelength of 450 nm.

### Saline and aprotinin blockage of furin- and trypsin-mediated FCS cleavage

His- and biotin-tagged peptides containing the wild-type and mutant (P681R) SARS-CoV-2-specific furin motif were added at a concentration of 5 ng/well (prepared with PBS) to the blocked (2% BSA) Ni-NTA-coated wells of the 8-well microplate strips and incubated for 1 h at RT. Meanwhile, the furin or trypsin diluted in PC solution was incubated with saline or aprotinin at different concentrations for 10 min. After washing the strips for 3 times with PBS-T, the enzyme solutions pre-incubated with NaCl (Sigma) or aprotinin (Sigma) at different concentrations were transferred into the strip wells and incubated for 25 min at 37°C. After washing for 4 times, streptavidin-HRP (100 μl, 1:5000 dilution in PBS-T) was added and incubated for 30 min at RT. After washing for 4 times again, 100 μl of TMB solution were added per well and blue color development was monitored for 2-10 min. The reaction was stopped with an equal volume of 1 M HCl and the optical density was measured with a microplate reader at a wavelength of 450 nm.

### Saline and aprotinin based inhibition of nasal protease-mediated FCS cleavage

His- and biotin-tagged peptides containing the wild-type and mutant (P681R) SARS-CoV-2-specific furin motif were added at a concentration of 5 ng/well (prepared with PBS) to the blocked (2% BSA) Ni-NTA-coated wells of the 8-well microplate strips and incubated for 1 h at RT. After washing the strips for 3 times with PBS-T, the nasal swab sample supernatant was incubated with aprotinin at different concentration for 10 min. The sample solutions were then transferred into the strip wells and incubated for 25 min at 37°C. After washing for 4 times, streptavidin-HRP (100 μl, 1:5000 dilution in PBS-T) was added and incubated for 30 min at RT. After washing for 4 times again, 100 μl of TMB solution were added per well and blue color development was monitored for 2-10 min. The reaction was stopped with an equal volume of 1 M HCl and the optical density was measured with a microplate reader at a wavelength of 450 nm.

### Statistical analysis

Data are presented as means ± SD. The cleavage percentage of enzyme-treated samples incubated with or without saline and aprotinin was calculated as follows:

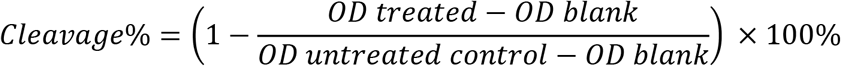

## Results

### Cleavage of both wild-type and mutant (P681R) FCS by furin and trypsin

The sequences of the synthesized peptides, as shown in the table 1, are based on the complete SARS-CoV-2 furin cleavage site characterized as a 20 amino acid motif (Fig 1).

**Table 1.**
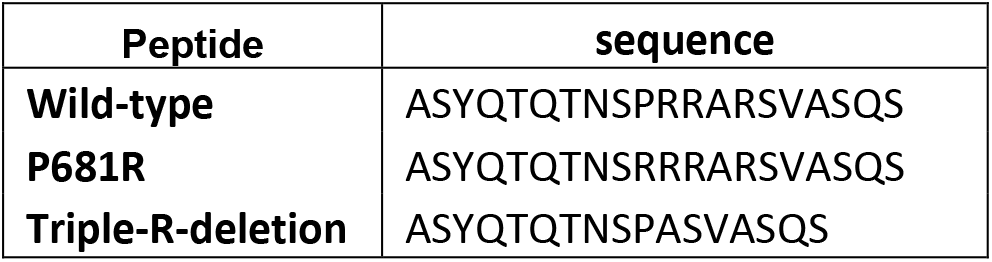
The list and sequence of the SARS-CoV-2 wild-type and mutant FCS peptides.

**Fig. 1.**
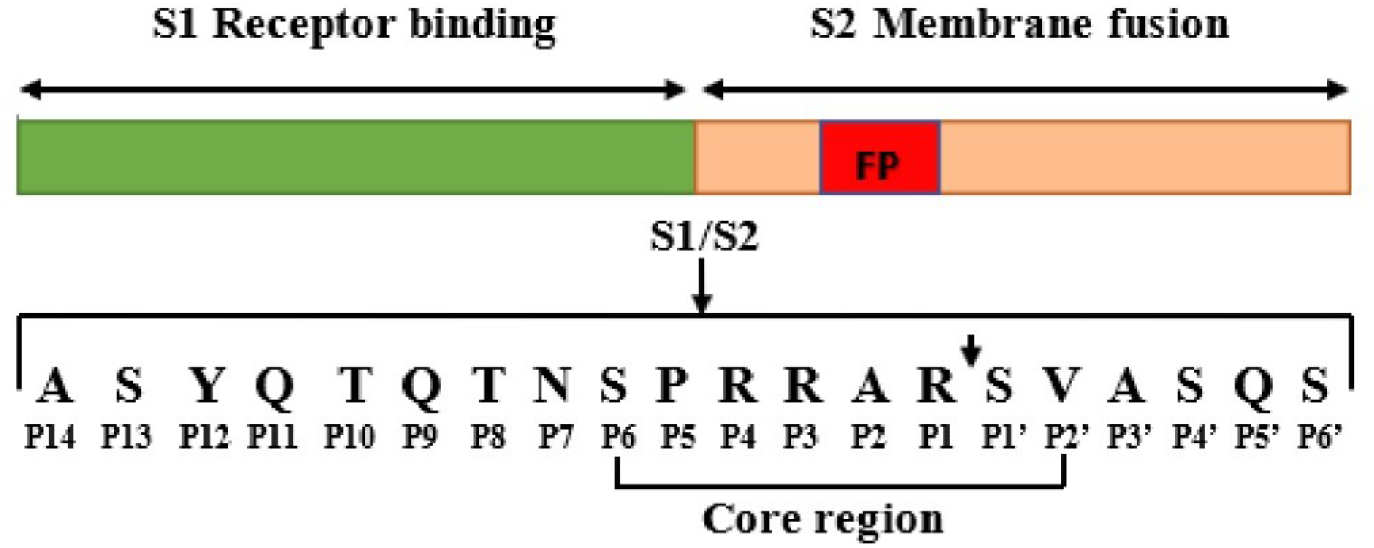
Schematic of the SARS-CoV-2 spike protein, and the furin cleavage site at the boundary between the S1 and S2 subunits.

The dual-tagged peptides with polyhistidine at the N-terminal and biotin at the C-terminal bind onto microplate wells through histidine/Ni-NTA at its N-terminal. The cleavage of the peptides in the FCS core region will remove the C-terminal part of the peptides after washing, which causes the decrease in signal generated by avidin/biotin binding after adding streptavidin-HRP. As shown in Fig 2A and 2B, the furin enzyme cleaved the wild-type and P681R mutant FCS, but not triple R-deleted FCS in a concentration dependent manner. Serine protease trypsin can cleave both wild-type and P618R FCS with higher cleavage percentage even at 10 ng of the concentration. However, the triple R-deleted FCS peptide was also not cleaved by trypsin. These results are consistent with our previously reported data [11]. And thus we focused on the wild-type and P681R types in the next studies.

**Fig. 2.**
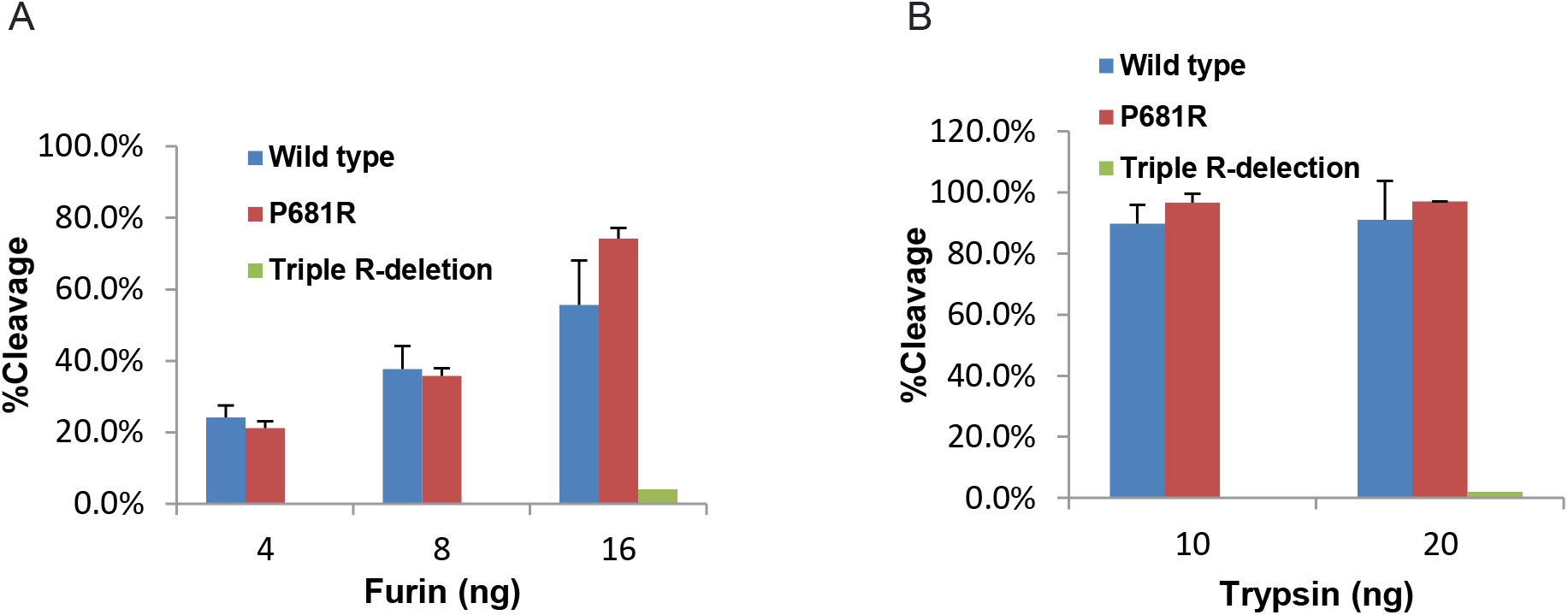
Cleavage of SARS-CoV-2 wild-type, P681R and triple R-deletion mutant FCS by serine proteases at different concentrations. A). furin; B). trypsin.

### Cleavage of both wild-type and mutant (P681R) FCS by nasal swab sample

We also tested the FCS cleavage of these peptides by nasal swab samples. Endogenous proteases from the nasal mucosa and exogenous proteases from the bacterial flora of the nasal cavity can be captured by nasal swab. As shown in Fig 3, both wild-type and P681R FCS peptides were cleaved by nasal swab solution in a concentration-dependent manner. At 25 μl solution volume that is only 10% of the collected sample supernatant volume, more than 80% of FCS peptides were cleaved.

**Fig 3.**
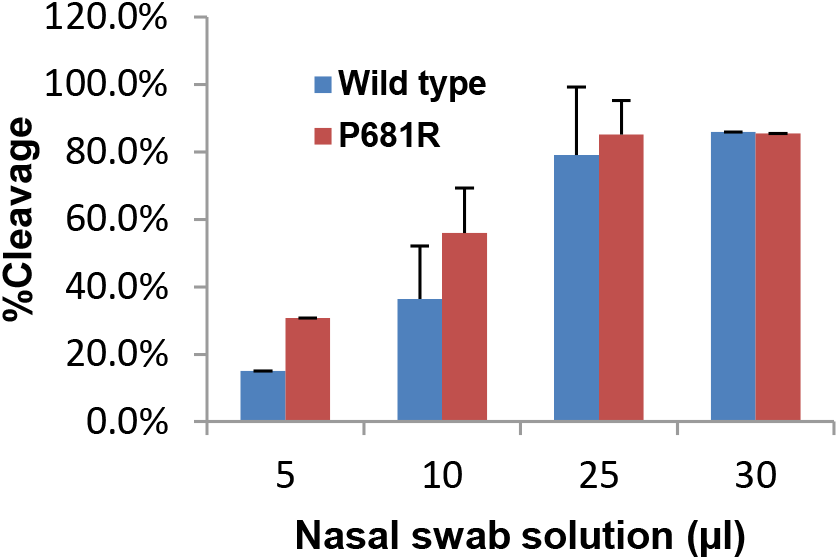
Cleavage of SARS-CoV-2 wild-type and P681R mutant FCS by nasal swab sample at different amount.

### Hypertonic saline and aprotinin block furin- and trypsin-induced cleavage of both wild-type and mutant (P681R) FCS

We further examined the effects of saline and aprotinin on FCS blockage of these peptides. Dual His- and biotin-tagged FCS peptides were bound to Ni-coated microplate wells. The wells were then exposure to furin or trypsin solution which was pre-incubated with saline or aprotinin at different concentrations. As shown in Fig. 4A, at a physiological concentration (0.9%), saline did not show the inhibitory effect on furin enzyme-caused cleavage of the FCS peptides. Higher saline concentrations effectively blocked the cleavage of both wild-type and P681R FCS caused by furin enzyme. At 3% of the concentration, around 80% of the furin cleavage effect was reduced in wild-type and 70% decreased in P681R type, respectively. In contrast, trypsin-caused cleavage of wild-type and P681R FCS was only slightly inhibited (< 10%) by saline even at hypertonic concentrations (Fig 4B). The aprotinin exhibited a dose-dependent inhibition of furin- and trypsin-induced cleavage of the wild-type and P681R FCS (Fig 4C and 4D). At 2 μg per well (around 6 μM), approximately 65-73% furin cleavage and 85-97% trypsin cleavage were reduced by aprotinin, respectively.

**Fig 4.**
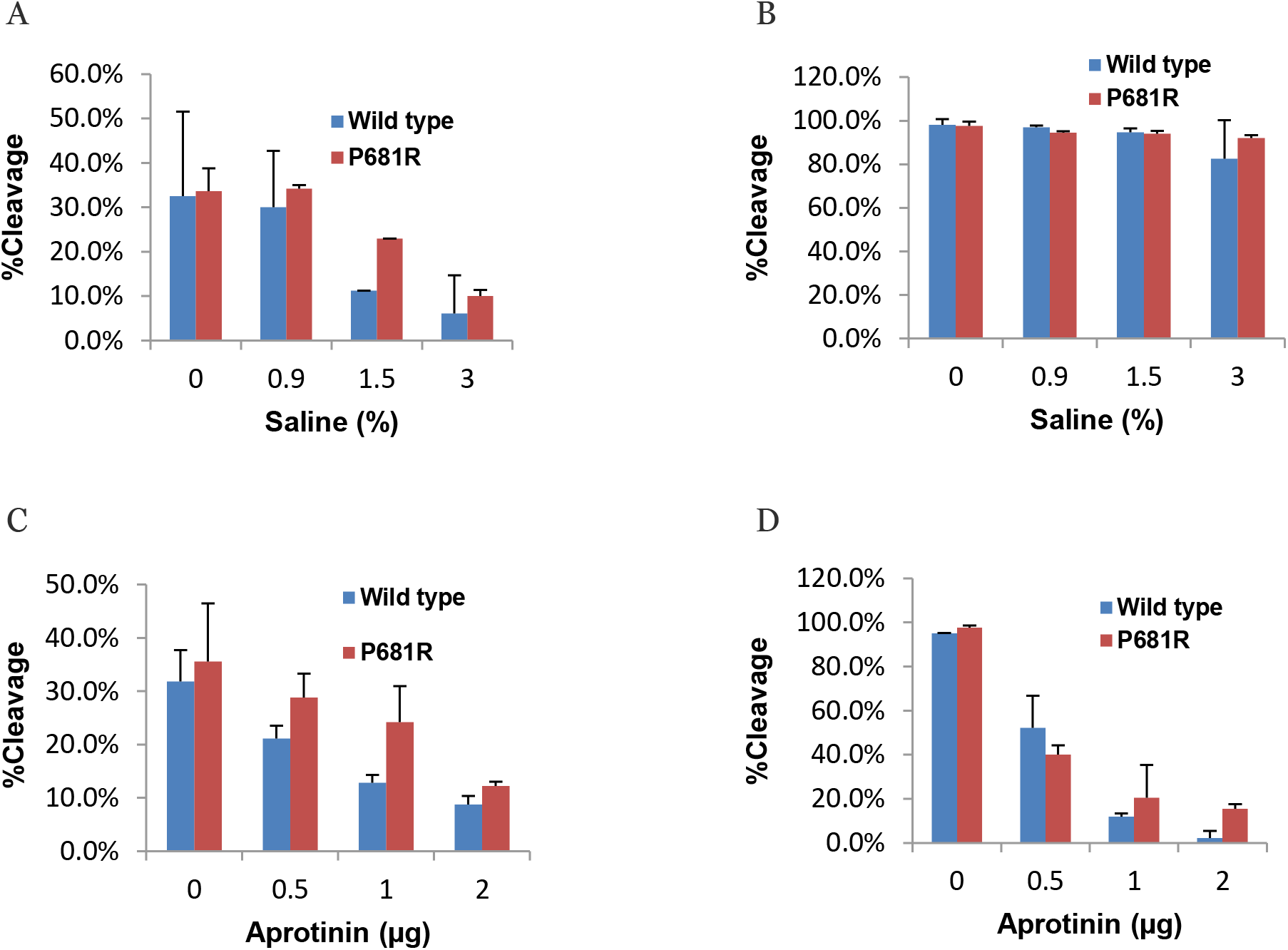
Blockage of SARS-CoV-2 wild-type and P681R mutant FCS cleavage caused by furin and trypsin. A) Saline blockage on furin (8 ng/well) cleavage; B). Saline blockage on trypsin (10 ng/well) cleavage; C). Aprotinin blockage on furin (8 ng/well) cleavage; D). Aprotinin blockage on trypsin (10 ng/well) cleavage.

### Hypertonic saline and aprotinin block nasal swab sample-caused cleavage of both wild-type and mutant (P681R) FCS by inhibition of nasal protease activity

Inhibition of nasal swab sample-caused cleavage of the peptide FCS by saline and aprotinin was next examined. As shown in Fig 5, in wild-type, the cleavage was reduced from 60.7% to 22% by hypertonic saline (3%) and from 60.7% to 41.5% by 2 μg of aprotinin, while in P81R mutant type, the cleavage was only decreased to 51.7% from 69.5% by hypertonic saline and to 63.5% form 69.5% by aprotinin. However, the reduction of cleavage was significant when combining hypertonic saline and aprotinin. In wild-type, nasal swab sample-caused cleavage was reduced to 0.5% from 60.7% and in P681R mutant type, the cleavage was decreased to 10% from 69.5%.

**Fig. 5.**
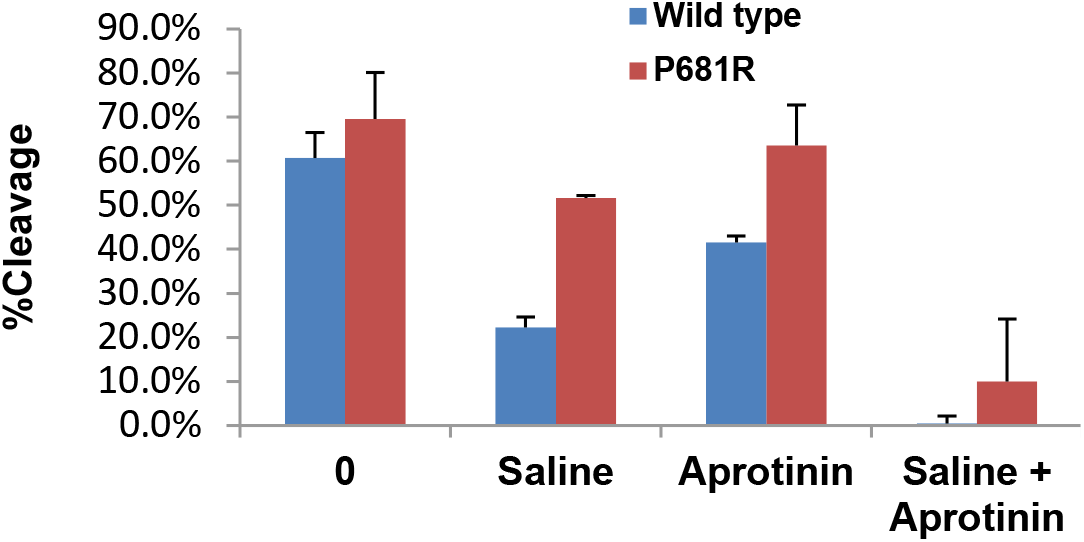
Blockage of SARS-CoV-2 wild-type and P681R mutant FCS cleavage caused by nasal swab samples. Saline alone: 3%; Aprotinin alone: 2 μg/well; Saline and aprotinin combination: 3% + 2 μg.

## Discussion

In this study, we utilized synthesized peptides containing wild-type or P681R mutant FCS to examine the effect of hypertonic saline and serine proteinase inhibitor aprotinin on blockage of SARS-CoV-2 specific furin site cleavage. Our previous study shows both SARS-CoV-2 spike protein and synthesized wild-type FCS peptide can be bound and immunocaptured by SARS-CoV-2-specific furin site blocking antibody in a concentration-dependent manner [11]. And both the synthesized peptide and full-length S-protein were cleaved to a similar level by recombinant furin protein. The results indicate that the synthesized peptide is the same as biological S protein as a SARS-CoV-2-specific FCS cleavage assay substrate.

It was demonstrated that saline results in a dose-dependent inhibition of replication of a range of DNA and RNA viruses, including the human coronavirus 229E (HCoV-229E) [12]. Hypertonic saline nasal irrigation and gargling was also found to reduce viral upper respiratory tract infection in a clinical trial [13, 14]. Recently, hypertonic saline (1.5% NaCl) was shown to significantly inhibit replication of SARS-CoV-2 in cultured human lung and kidney epithelial cells. The major mechanism is possibly through sodium ion-mediated intracellular low energy states [15]. In this study, we observed that hypertonic saline can significantly block SARS-CoV-2-specific FCS cleavage caused by furin but not by trypsin. As S protein cleavage plays a critical role in SARS-CoV-2 activation before entry into and exiting from cells, the inhibitory effect of saline on SARS-CoV-2 replication could be at least partly due to saline blockage of furin-mediated cleavage. Aprotinin is a broad serine protease inhibitor and approved by the FDA for clinical use for reducing bleeding during complex surgery [16]. However, it showed significant antiviral effects against H1N1 influenza infections both in vitro and in vivo by blocking HA cleavage and activation and is also approved in Russia for use as a nasal aerosol for the treatment of influenza [17]. Bojkova et al recently reported that aprotinin is able to inhibit SARS-CoV-2 replication to block SARS-CoV-2-induced cytopathogenic effect (CPE) formation at therapeutically achievable concentrations [18]. Our study shows that the aprotinin effect at 3-6 μM level is remarkable in decreasing trypsin-mediated SARS-CoV-2 FCS cleavage but only moderate in inhibiting furin-based FCS cleavage.

We observed that higher cleavage percentages were achieved with both furin and trypsin for P681R FCS peptide than for wild-type one. It is consistent with that the Delta variant’s P681R allows the cleavage to occur more easily than in the wild-type SARS-CoV-2 strain, as Delta variant has a more efficient furin cleavage site [19,20]. However, the cleavage of P681R can still be inhibited by hypertonic saline or aprotinin at an appropriate concentration, even if such inhibition was relatively less in P681R FCS compared to wild-type. An interest observation in this study is that the combination of hypertonic saline and aprotinin achieved nearly complete inhibition of FCS cleavage in both wild-type and P681R mutant with use of nasal swab samples, although hypertonic saline or aprotinin alone only has moderate and mild inhibitory effect on both wild-type and P618R mutant, respectively. This additive effect would suggest the importance of the combination of hypertonic saline and aprotinin in practical use for SARS-CoV-2-specific FCS cleavage inhibition, as the nasal proteases can be from both host cells and bacteria in the nasal cavity. These proteases relevant to FCS cleavage could include both furin/furin-like proprotein convertases targeting furin cleavage motif and trypsin/trypsin-like serine proteases targeting mono or multiple arginine in FCS, which may be with different sensitivity to hypertonic saline and aprotinin.

It is generally accepted that the nasal epithelium is the initial source of SARS-CoV-2 infection and proliferation [21, 22], where nasal protease-mediated S protein cleavage causes vial activation and replication [23]. In the absence of effective drugs and decrease in the vaccine effect against Delta variant, topical application of hypertonic saline combining with aprotinin such as nasal drop or aerosol may represent simple, economical, practically feasible approach, which would be greatly helpful in locally controlling viral activation and entry into cells to replicate, thereby reducing/preventing SARS-CoV-2 infection and avoiding covid-19 progress to a severe and systemic disease.

## Competing Interest Statement

The authors have declared no competing interest.

## funding

This work was supported by Epigentek Group, Inc.

